# Physiology and Mathematical Modeling of Immobilized *Saccharomyces* spp. in Beer Fermentation

**DOI:** 10.1101/2022.12.17.520861

**Authors:** Thiago M. de Araujo, Marcel M. L. da Cunha, Marcelo C. Barga, Bianca E. Della-Bianca, Thiago O. Basso

## Abstract

There is an ever-increasing demand for reduction of unit operations and a growing interest in the physiology of yeasts used in beer fermentation. In this context, cell immobilization is an interesting alternative, since it reduces steps to separate biomass from fermented broth. Yet, physiological alterations in yeast metabolism caused by immobilization are still to be fully described. Thus, the main objective of this work was to evaluate the physiology of three brewer’s *S. cerevisiae* yeast strains (SY025, SY067 and SY001) immobilized on a porous cellulose-based support. Batch fermentations in malt extract 12 °P were carried out for all strains both in free and immobilized forms in order to compare kinetic parameters obtained from distinct process conditions. Mathematical modeling was performed following two viewpoints: modeling of fermentation kinetics by parameter estimation from experimental data and application of a reaction-diffusion model for estimation of substrate concentration gradient inside the immobilization support. Moreover, fermentations with different initial substrate and biomass concentrations were carried out using strain SY025, aiming to evaluate their influence over flavor compounds, using statistical models. Compared to free cells, immobilized yeasts showed both higher glycerol yield (SY025, 40%; SY067, 53%; SY001, 19%) and biomass yield in the system (SY025, 67%; SY067, 78%; SY001, 56%). On the other hand, free cells presented higher ethanol yields when compared to immobilized ones (SY025, 9%; SY067, 9%; and SY001, 13%). According to the model developed, a substrate gradient inside the support was predicted, but with low mass transfer limitations.

**KEY POINTS:**

- Yeast immobilization not always hinder biomass growth, here it was stimulated.
- A classic kinetic model describes accurately immobilized yeast fermentations.
- Physiology changes occur in immobilization even with low mass transfer limitations.

## 1. INTRODUCTION

Cell immobilization is their confinement within a delimited region by means of physical and/or chemical mechanisms, in general using some solid support called immobilization matrix (Kourkoutas et al. 2004; Moreno-García et al. 2018; Pilkington et al. 1998). This technology started in the 1960s by adapting enzyme immobilization techniques to living cells (Karel et al. 1985; Nedović et al. 2015).

In industrial biotechnology, cell immobilization offers several advantages in comparison to traditional fermentations with suspended cells, such as higher cell density, productivity and cell resistance to shear stress, and lower bioreactor volume and operation costs (Kosseva 2011; Rouf et al. 2017). Moreover, this technology results in reduced contamination risk (assuming sterile conditions in the immobilization step), due to high cell density (Nedovic et al. 2011), and elevated dilution rates during continuous processes. Easy separation of biomass from the final product and cell reuse are also important features of this technology (Boulton and Quain 2001; Kourkoutas et al. 2004).

However, there are some challenges and drawbacks, such as mass transfer limitations (Norton and D’Amore 1994; Radovich 1985), clogging, fragile supports disruption and expressive physiological alterations that may result in esters and acetaldehyde overproduction above the sensorial threshold, which affects final product quality, in case of beer fermentation (Boulton and Quain 2001). Furthermore, scaling-up and standardization of both immobilization support production and immobilized cell process operation are very complex (Kourkoutas et al. 2004; Norton and D’Amore 1994).

In brewery fermentation, yeast immobilization had been widely investigated from 1970s to 1990s to develop a continuous process. It was believed this technology would revolutionize the brewing industry by increasing productivities (Virkajärvi and Linko 1999). However, decades of research demonstrated that immobilization procedures, process conditions and bioreactor configuration negatively affected beer quality due to direct impact on the physiology and metabolism of cells; flavor profiles in the final product rarely matched those achieved by traditional batch fermentation (Willaert and Nedovic 2006). Moreover, inherent risk of contamination in continuous systems was also a concern. The focus was on industrial scale production of lager beer (Nedović et al. 2015; Virkajärvi and Pohjala 2000; Yamauchi et al. 1994), in which any variation from a strict standard quality control is undesired, and so the immobilization technology became not commercially attractive.

During the last 20 years, however, there was an important expansion on the beer market, especially regarding beer styles variation and new flavor profiles. Even the big market players have incorporated craft beer styles to their scope to remain competitive (Morgan et al. 2020). And this rising market is one of the main factors to reawaken interest on brewery fermentation using immobilized yeast in batch process, in which physiological alterations might be very welcome, and not necessarily undesired due to the diversity of beer styles produced by craft breweries (Araujo et al. 2021).

Even though much has been investigated about cell immobilization, two main topics remain open: efficient scale-up of industrial process and full comprehension of the mechanisms that lead to alterations on cell physiology (Kourkoutas et al. 2004; Norton and D’Amore 1994). Therefore, our objective was to investigate the physiological changes of three brewer’s yeast strains, immobilized to a porous solid cellulose-based support (namely ImoYeast, developed by Bio4 Soluções Biotecnológicas Ltda.), by analyzing and modeling their fermentation kinetics. Finally, a design of experiments was developed to evaluate the influence of initial concentrations of biomass (X_0_) and substrate (S_0_) over flavor compounds.

## 2. MATERIALS AND METHODS

### 2.1 Yeast strains

All yeast strains used in this study were provided by Bio4 Soluções Biotecnológicas Ltda.: Ale (American Ale - SY025, *Saccharomyces cerevisiae*); Weiss (Belgian Wit - SY067, *Saccharomyces cerevisiae*); and Lager (Pilsner Lager -SY001, *Saccharomyces pastorianus*). Stocks of all strains were kept at -80 °C in 20% (v/v) glycerol.

### 2.2 Yeast immobilization

ImoYeast, also manufactured and provided by Bio4, was used as immobilization matrix. It is a cellulose-based, round-shaped, porous solid able to absorb biomass suspensions, with diameter of ca. 7.5 mm.

A 20 mL yeast suspension grown in YPM 12 °P was centrifuged by 15 min at 10,000 g in a refrigerated centrifuge CR 21GIII (Hitachi, Japan). Next, supernatant was discarded and to the left biomass it was added 50 mL sodium alginate solution 3% (m/ v) and homogenized.

This new suspension was then absorbed by dried ImoYeast particles at proportion 18 g support to 20 mL suspension. The system was submerged in 250 mL calcium chloride suspension 2% (m/ v) by 30 min. ImoYeast matrix containing biomass was coated by 50 mL sodium alginate 3% (m/ v) by 10 s. Finally, particles were submerged in 250 mL calcium chloride solution 2% (m/ v) by 30 min. After cell immobilization, the particles were not washed.

### 2.3 Fermentation assays

Fermentation kinetics were carried out in triplicate for all strains, in both suspended and immobilized configurations.

All experiments were performed in static batch mode in 500 mL Erlenmeyer flasks containing 200 mL malt wort 12 °P (pilsen malt wort Grano Mestre, Iomerê, lot 0119) and sealed by silicone stoppers equipped with one-way filters for gas release. Different initial biomass concentrations and temperatures were used for each strain, according to industrial practice as follows: SY025 – 9.0×10^6^ cells/mL at 18 °C; SY067 – 1.2×10^7^ cells/mL at 20 °C; and SY001 – 1.8×10^7^ cells/mL at 11 °C (White and Zainasheff 2010). Fermentations took 4-5 days and daily samples were taken for quantification of biomass, substrate and product concentrations.

#### 2.3.1 Analytical methods

Biomass concentration (cells/mL) was quantified by Neubauer chamber counting using 1 mL of methylene blue 0.2% in 1 mL of sample, homogenized by 1 min. Sample dilution ranged from 2x to 100x, aiming each Neubauer chamber’s quadrant to contain up to 30 cells. Total number of free cells was calculated by multiplying biomass concentration and the culture volume in the flask.

Immobilized biomass quantification needed additional steps, as follows. Initially, ImoYeast particles (1-2 units) were withdrawn from culture and then weighed for determining wet weight. Dry weight was obtained by multiplying wet weight by 0.27, a factor previously characterized for ImoYeast particles dry weight (data not shown). Particles were then resuspended in 2 mL sodium phosphate 5% solution (m/ v) for releasing the cells, which were counted by the Neubauer chamber method (Pilkington et al. 1998).

For both suspended and immobilized cells, cell number data was converted into total biomass concentration using correlations obtained to each strain (data not shown).

Substrates and products were quantified by HPLC Prominence (Shimadzu), using: ion exchange column Aminex HPX-87H (Bio-RAD) at 60 °C, sulfuric acid solution 5 mM as mobile phase and flow at 0.6 mL/ min; and IR and UV detectors at 50 °C.

For flavor profiling, beer was collected at the end of fermentation, centrifuged at 10,000 g by 10 min in a refrigerated centrifuge CR 21GIII (Hitachi, Japan) and the supernatant was frozen. After thawing, the samples were analyzed by GC-FID using an Elite WAX Polyethylene glycol column (length 60 m, ID 0.25 mm, film thickness 0.25 μm). Operation system parameters were: 1 μL automatic injection via auto sampler; injector temperature 220 °C; detector temperature 250 °C; and initial kiln temperature 40 °C, ramping up by 5 °C/ min until 70 °C, isotherm by 7.4 min and ramping up by 25 °C/ min until 90 °C, and then isotherm by 5.8 min.

Calibration curves were elaborated using GC standards (Merck) for 1-propanol, 2-methyl propanol (isobutyl alcohol), 3-methyl butanol (isoamyl alcohol), ethyl acetate and acetaldehyde, with 1-butanol 50 mg/ L as internal standard.

#### 2.3.2 Scanning Electron Microscopy (SEM)

Samples of ImoYeast particles were periodically taken during fermentation (approximately 0 h, 69 h and 117 h), then suspended in 2 mL reverse osmosis water and refrigerated.

Particles were fixed in 2.5% glutaraldehyde and 4% paraformaldehyde in 0.1 M cacodylate buffer pH 7.2 for at least 1 hour, washed in the same buffer. Next, the particles were dehydrated in 30, 50, 70, 90 and 100% ethanol and critical point-dried with CO_2_ (Cunha et al. 2005).

To observe the distribution of yeast inside ImoYeast particles, samples were individually frozen in liquid nitrogen and fractured cross-sectionally with a scalpel submerged in liquid nitrogen. Finally, using a desiccator connected to a vacuum pump, samples were dried, mounted in stubs with conductive double sided carbon tape and sputter coated with a thin layer of gold in a Denton Vacuum Desk V (Denton Vacuum Inc., USA) apparatus for observation in a TESCAN VEGA 3 (Brno, Czech Republic) scanning electron microscope.

#### 2.3.3 Estimation of kinetic parameters

Mass balance equations were used for suspended (Equations 1 to 4) and immobilized (Equations 5 to 9) cell systems to describe biomass growth, substrate consumption and products formation kinetics.

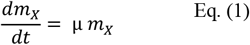

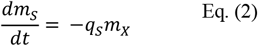

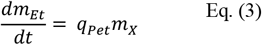

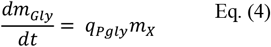

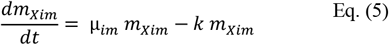

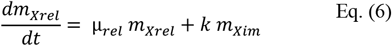

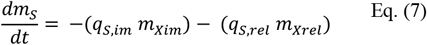

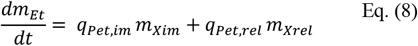

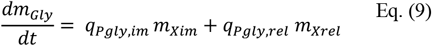

Kinetic equations are listed for suspended (Equations 10 to 13) and immobilized (Equations 14 to 21) cells as follows.

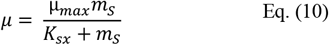

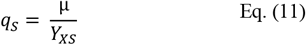

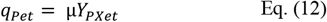

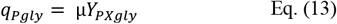

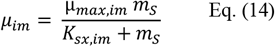

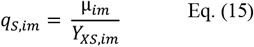

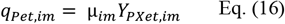

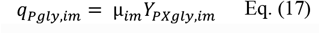

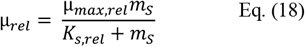

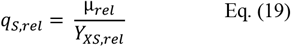

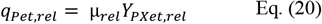

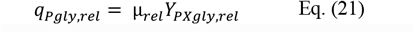

A Michaelis-Menten-type behavior was assumed for the kinetics of cell release from the support, as shown in Equation 22.

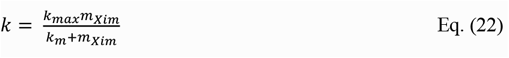

Models were implemented in Python programming language (Spyder, version 3.8) using the Nelder-Mead parameter estimation technique (Barati 2011). Initial conditions were obtained from experimental data.

#### 2.3.4 Reaction-diffusion model

Following Doran (2013), there were assumptions to the model: the ImoYeast support is considered a perfect sphere; the particle is homogeneous, isothermal and isotropic; mass transfer occurs by diffusion only (Fick’s law); substrate variation occurs in one single spatial variable (radius of the particle); substrate consumption rate is governed by Monod kinetics; and the control volume operates in steady state.

Substrate mass balance in a spherical particle was developed by the shell method (Doran 2013; Engasser and Horvath 1973; Golman 2016; Shuler and Kargi 2002) in the control volume and it is expressed in its dimensionless form by Equation 23.

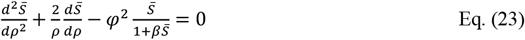

With boundary conditions defined by Equations 24 and 25.

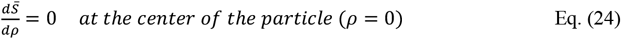

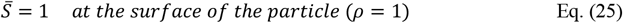

All the variables present in the model are described as follows, by Equations 26 to 29.

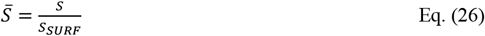

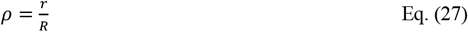

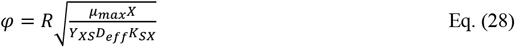

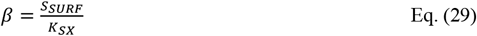

D_eff_ was calculated according to the empirical relation proposed by Axelsson and Persson (1988), using standard sucrose diffusivity in calcium alginate 2% (m/ v) at 25 °C (D_SUC_ = 1.73×10^−6^ m^2^/ h) (Doran 2013). It is presented by Equation 30.

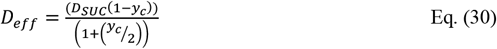

Where the biomass fraction inside the solid particle (y_c_) is given by Equation 31.

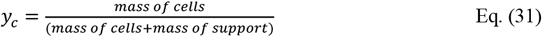

The model solution was carried out based in Golman (2016), using the finite differences numerical method. Simulations were performed with Python on Jupyter Notebook environment.

### 2.4 Variation of initial biomass and substrate concentrations

A central composite design experiment was performed (Table 1) to evaluate the impact of initial biomass (X_0_) and substrate (S_0_) concentrations over the production of flavor compounds by strain SY025 at 18 °C.

**Table 1.**
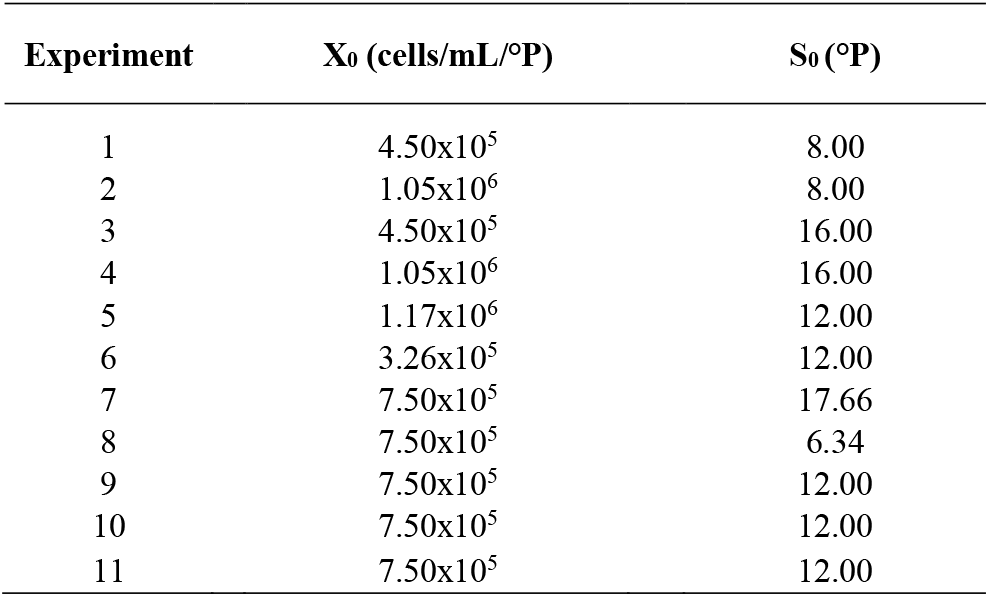
Calculated levels of initial biomass concentration (X_0_) and initial substrate concentration (S_0_) to each experiment.

| Experiment | $X_0$ (cells/mL/°P) | $S_0$ (°P) |
| --- | --- | --- |
| 1 | $4.50 \times 10^5$ | 8.00 |
| 2 | $1.05 \times 10^6$ | 8.00 |
| 3 | $4.50 \times 10^5$ | 16.00 |
| 4 | $1.05 \times 10^6$ | 16.00 |
| 5 | $1.17 \times 10^6$ | 12.00 |
| 6 | $3.26 \times 10^5$ | 12.00 |
| 7 | $7.50 \times 10^5$ | 17.66 |
| 8 | $7.50 \times 10^5$ | 6.34 |
| 9 | $7.50 \times 10^5$ | 12.00 |
| 10 | $7.50 \times 10^5$ | 12.00 |
| 11 | $7.50 \times 10^5$ | 12.00 |

All fermentations were conducted in static batch mode in 500 mL Erlenmeyer flasks sealed with cotton plugs and containing 200 mL malt wort (pilsen malt syrup Grano Mestre, Iomerê, lot 0320) by 8 days. Flavor compounds 1-propanol, 2-methyl propanol (isobutyl alcohol), 3-methyl butanol (isoamyl alcohol), ethyl acetate and acetaldehyde were quantified at the end of the fermentation.

Response Surface Methodology (RSM) was applied using Statistica to analyze the impact of the variables over the production of the quantified flavor-active compounds.

## 3. RESULTS

### 3.1 Fermentation kinetics data evidenced physiological alterations in immobilized yeast cells

From fermentation experiments with both immobilized and free cells, biomass growth data (Fig. 1) suggest cell growth stimulation in the immobilized condition. Immobilization efficiency was 70% (for strain SY025), 60% (SY067) and 90% (SY001) at the end of fermentation.

**Fig. 1.**
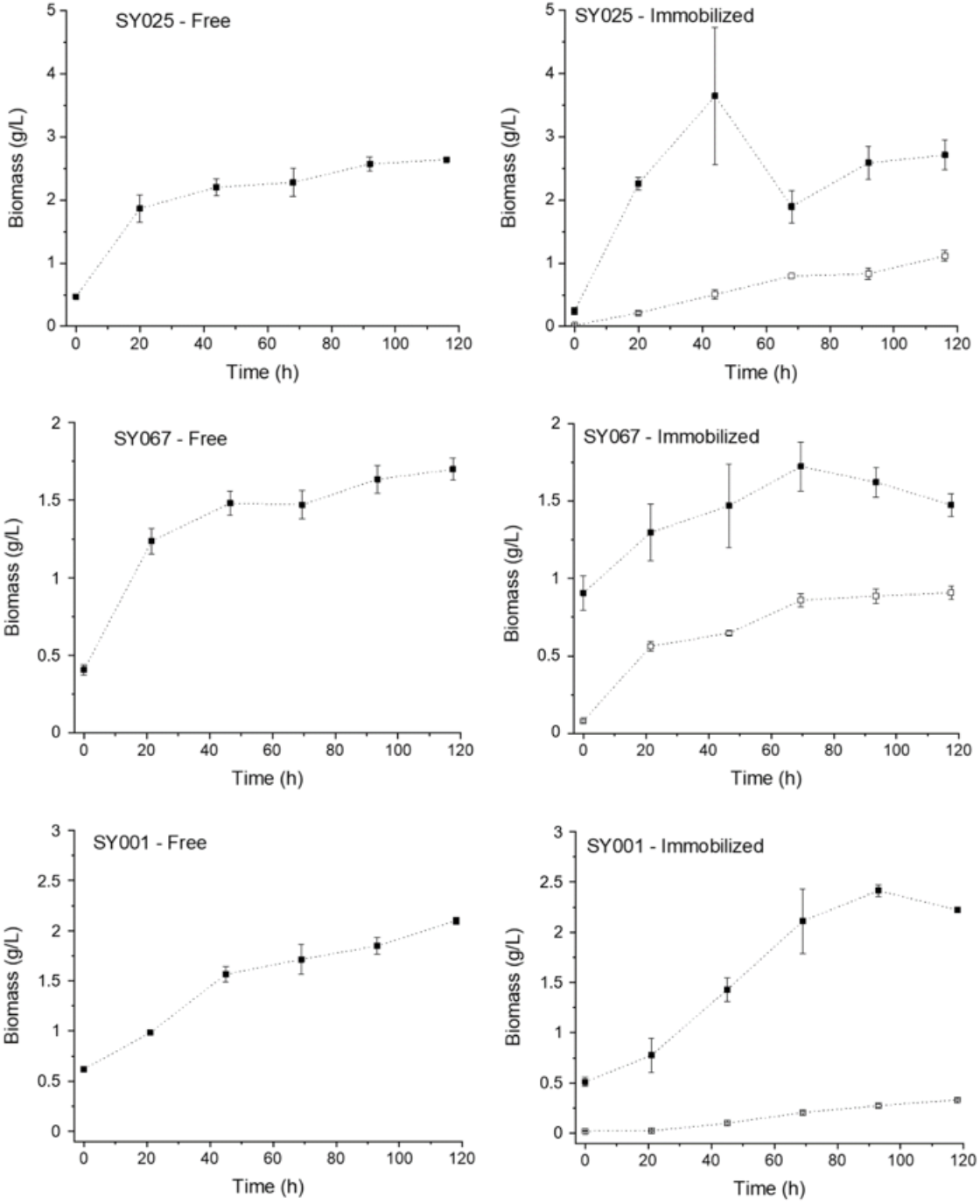
Biomass growth during beer fermentation using free or immobilized cells of different brewing yeast strains, in which: □ – yeast cells in suspension or released from ImoYeast support; and ▪ – immobilized yeast cells. Data and error bars represent mean values and standard error of three replicates, respectively.

For all strains evaluated, immobilized yeast systems produced more glycerol and less ethanol when compared to free cells (Fig. 2).

**Fig. 2.**
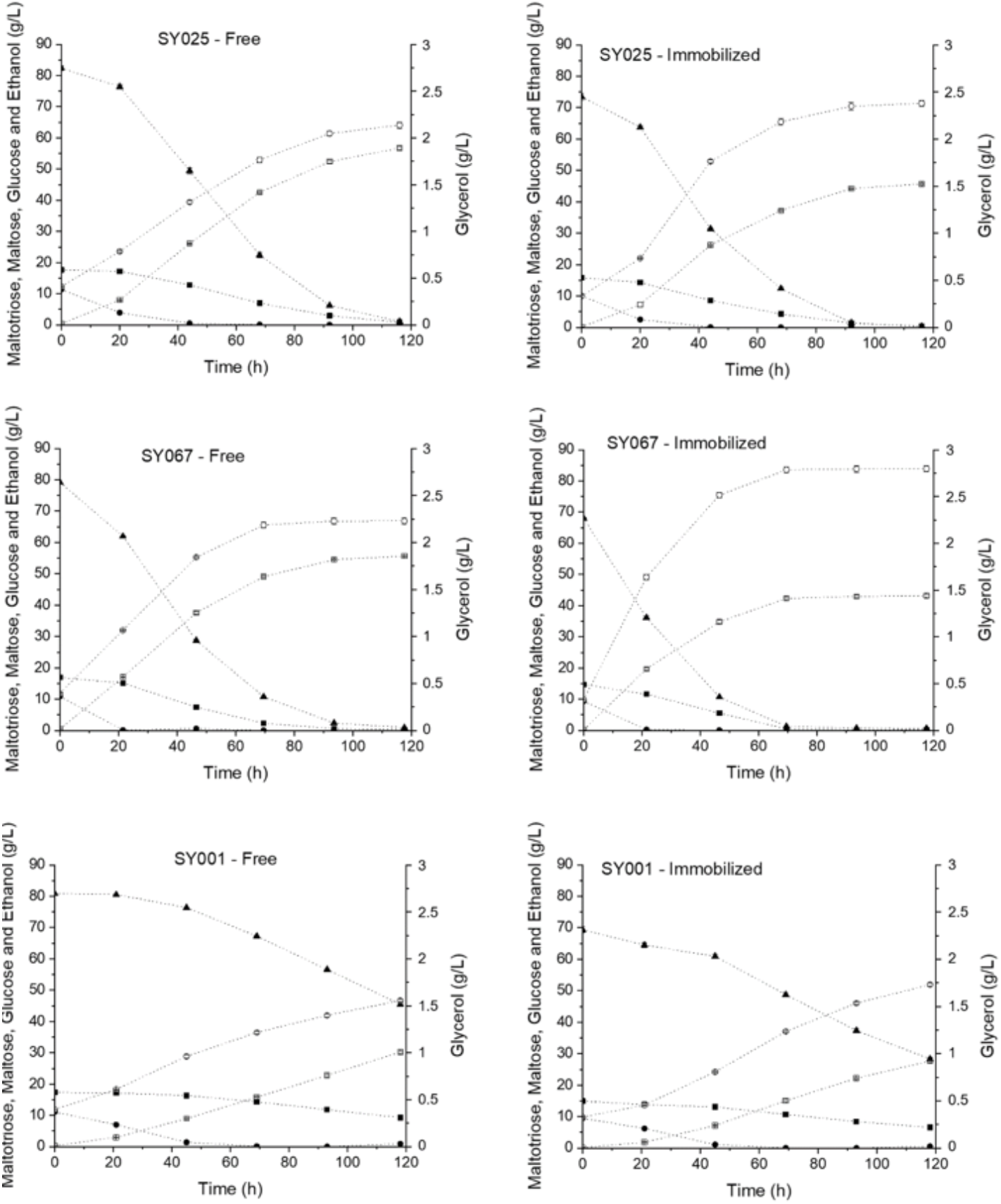
Substrate (▪ maltotriose, • glucose and ▴ maltose) consumption and product (□ ethanol and ○ glycerol) formation during beer fermentation with free or immobilized cells of strains SY025, SY067 and SY001. Data and error bars represent mean values and standard error of three replicates, respectively.

Global physiological parameters were obtained as presented by Table 2.

**Table 2.** Global physiological parameters obtained from fermentation kinetics for strains SY025, SY067 and SY001, both free and immobilized. Y_XS, im_= immobilized cells yield from substrate; Y_XS, TOTAL_= total biomass in the system yield from substrate; Y_PSet_= ethanol yield from substrate; Y_PSgly_ = glycerol yield from substrate; P_X, im_ = immobilized biomass mass productivity; P_X_ = total biomass mass productivity; P_Pet_ = ethanol mass productivity; and P_Pgly_ = glycerol mass productivity. Data and error represent mean values and standard error of three replicates, respectively. Total biomass in the system = immobilized + released biomass. *No statistically significant difference between free and immobilized cells.

| Global physiological parameters | SY025 |  | SY067 |  | SY001 |  |
| --- | --- | --- | --- | --- | --- | --- |
|  | Free | Immobilized | Free | Immobilized | Free | Immobilized |
| $Y_{XS, im}$ (g immobilized biomass/g substrate) | - | 0.016±0.001 | - | 0.007±0.001 | - | 0.033±0.003 |
| $Y_{XS, TOTAL}$ (g biomass/g substrate) | 0.015±0.001 | 0.025±0.001 | 0.009±0.001 | 0.016±0.001 | 0.025±0.000 | 0.039±0.003 |
| $Y_{PSet}$ (g ethanol/g substrate) | 0.509±0.000 | 0.464±0.004 | 0.516±0.000 | 0.471±0.004 | 0.553±0.000 | 0.489±0.008 |
| $Y_{PSgly}$ (g glycerol/g substrate) | 0.015±0.000 | 0.021±0.000 | 0.017±0.001 | 0.026±0.000 | 0.021±0.000 | 0.025±0.000 |
| $P_{X, im}$ (g immobilized biomass/h) | - | 0.021±0.002 | - | 0.005±0.002 | - | 0.014±0.001 |
| $P_X$ (g biomass/h) | 0.019±0.000 | 0.031±0.002 | 0.011±0.000* | 0.012±0.002* | 0.013±0.000 | 0.017±0.001 |
| $P_{Pet}$ (g ethanol/h) | 0.486±0.005 | 0.393±0.002 | 0.470±0.001 | 0.366±0.004 | 0.254±0.003 | 0.233±0.003 |
| $P_{Pgly}$ (g glycerol/h) | 0.015±0.000 | 0.018±0.000 | 0.016±0.000 | 0.021±0.000 | 0.010±0.000 | 0.012±0.000 |

Major flavor compounds were quantified at the end of the fermentation assays, for all strains (Fig. 3). Immobilized SY025 cells showed a distinct flavor profile than free cells of the same strain. Data suggest that immobilized SY025 metabolism was channeled into 1-propanol, ethyl acetate and acetaldehyde. Regarding strain SY067, immobilized cells presented lower production of 1-propanol and acetaldehyde and increased formation of 2-methyl propanol, 3-methyl-butanol and ethyl acetate when compared to free cells. Finally, in comparison to free cells, fermentation with immobilized SY001 strain resulted in lower concentrations of acetaldehyde and increased production of all other quantified flavor compounds.

**Fig. 3.**
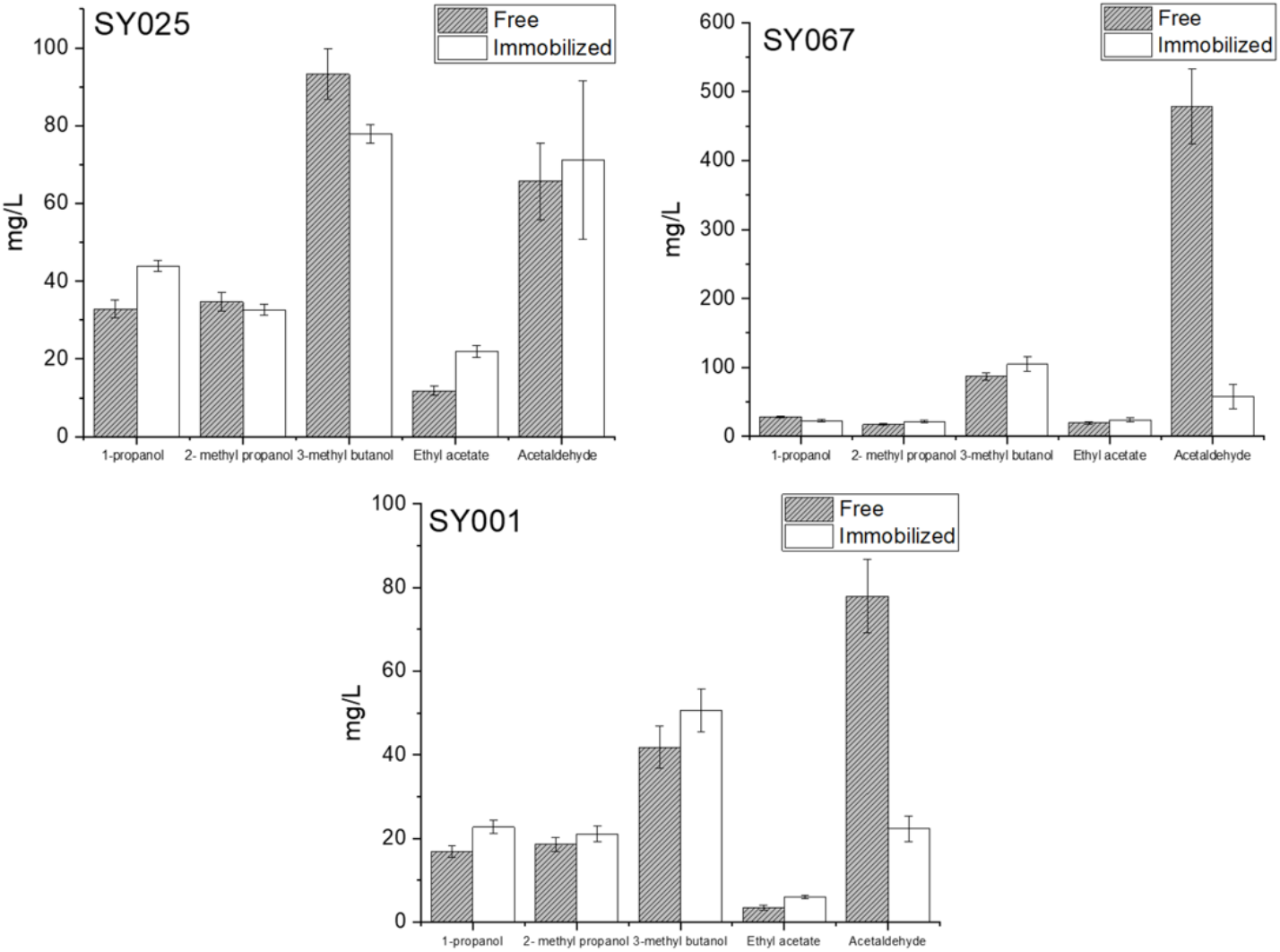
Flavor profiles obtained after 120 hours of beer fermentation for strains SY025, SY067 and SY001. Data and error bars represent mean values and standard error of three replicates.

### 3.2 Mathematical modeling – two distinct approaches indicate induced biomass growth of immobilized cells

The first approach in modeling aimed to determine kinetic parameters by mathematical optimization and then directly compare estimated data with the experimental set (Fig. 4).

**Fig. 4.**
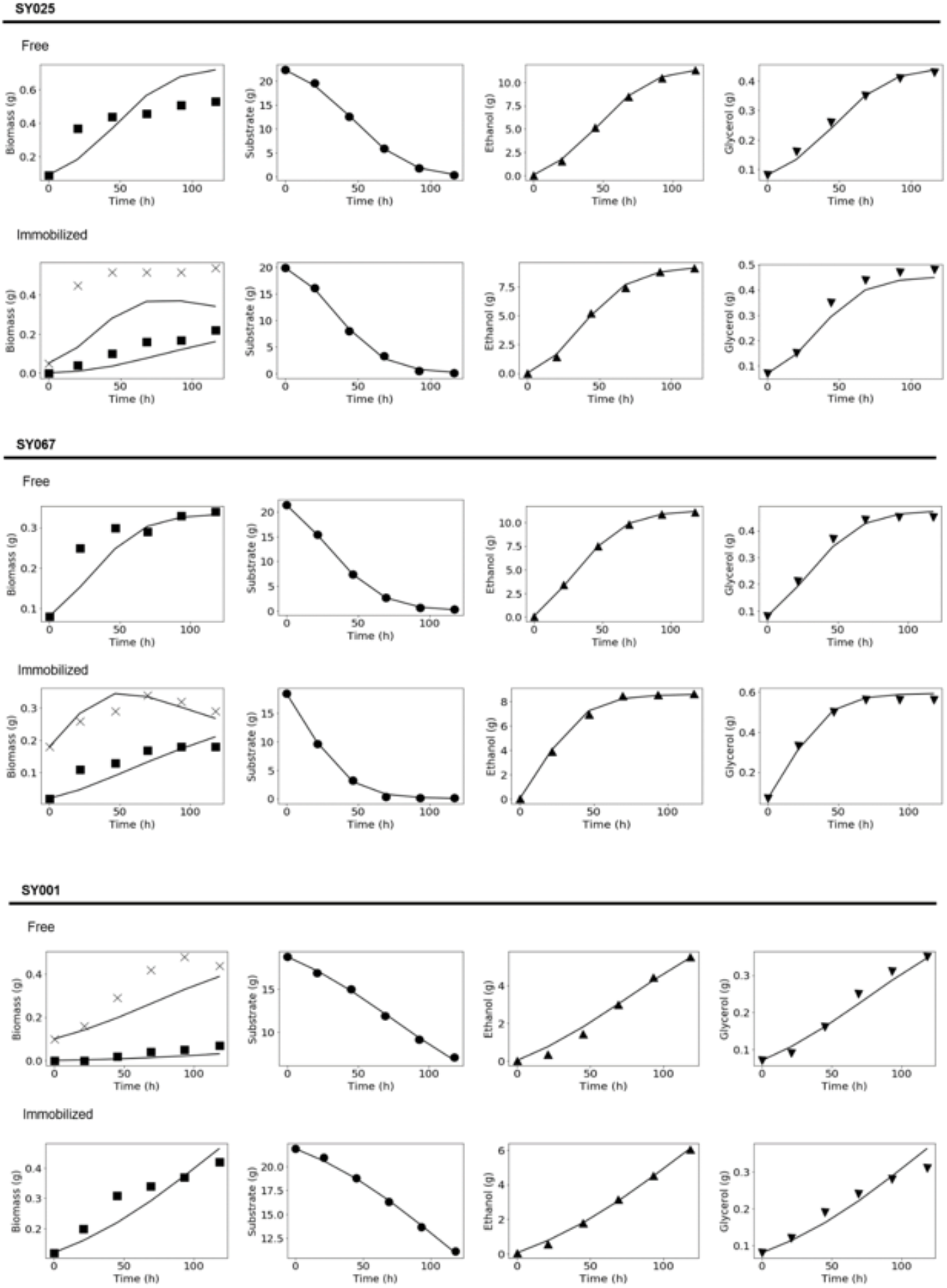
Experimental data x model, for strains SY025, SY067 and SY001 during fermentation kinetics assays, in which: ─ = model calculated values; X = immobilized biomass experimental data (g); ▪ = suspended/released biomass experimental data (g); • = substrate experimental data (g); ▴ = ethanol experimental data (g); and ▼ = glycerol experimental data (g).

For free SY025 cells, the Monod kinetic model was well adjusted to substrate, ethanol and glycerol experimental data. However, initial and final biomass values were under- and super-estimated, respectively (model residue of 0.43). Similar results were obtained for immobilized SY025 cells, but in this case biomass values were underestimated for the whole experiment, with residue of 0.89.

For strain SY067, modeling of the fermentation data for both free (model residue of 0.09) and immobilized (residue of 0.51) cells resulted in well-adjusted curves for biomass, substrate, ethanol and glycerol.

At last, for both free and immobilized SY001 cells the model was well adjusted to substrate and ethanol data, while biomass and glycerol presented less accurate results (model residues of 0.26 for free cells and 0.7 for immobilized cells).

Table 3 shows physiological parameters estimated by the model.

**Table 3.**
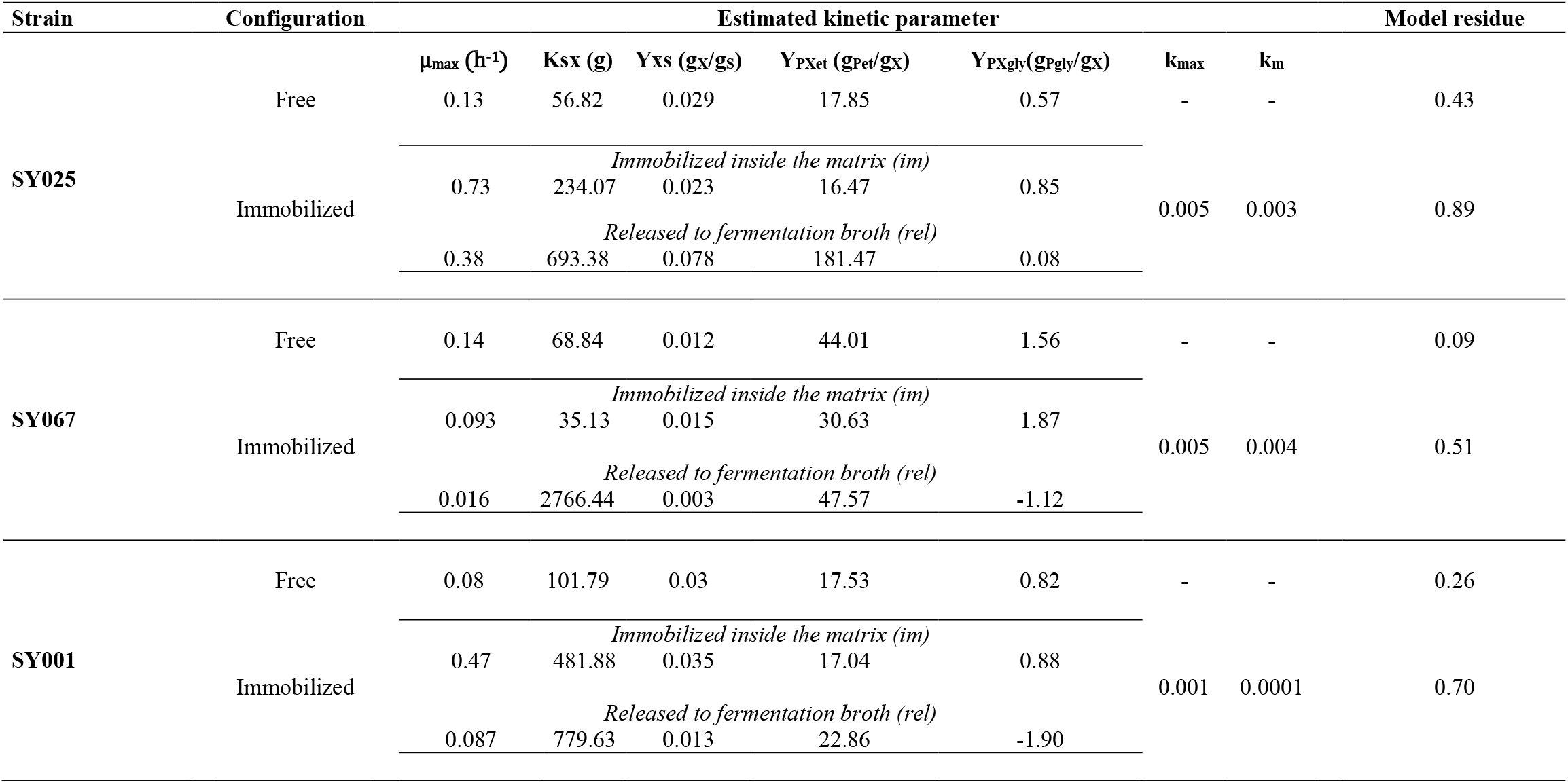
Kinetic parameters estimated by modeling for strains SY025, SY067 and SY001 during fermentation kinetics assays.

The second approach consisted in evaluating space distribution of substrate concentration within biocatalyst pellets. Since the model did not consider time variation (i.e., steady state), results are shown to each sampling time. Engineering parameters are also presented: Thiele modulus (φ), which defines a relation between mass transfer and reaction rates; effectiveness factor (η), a quantification of the support efficiency in comparison to a system with no mass transfer limitations; and saturation parameter (β), a ratio of substrate concentration at the solid surface and the saturation constant (Fig. 5).

**Fig. 5.**
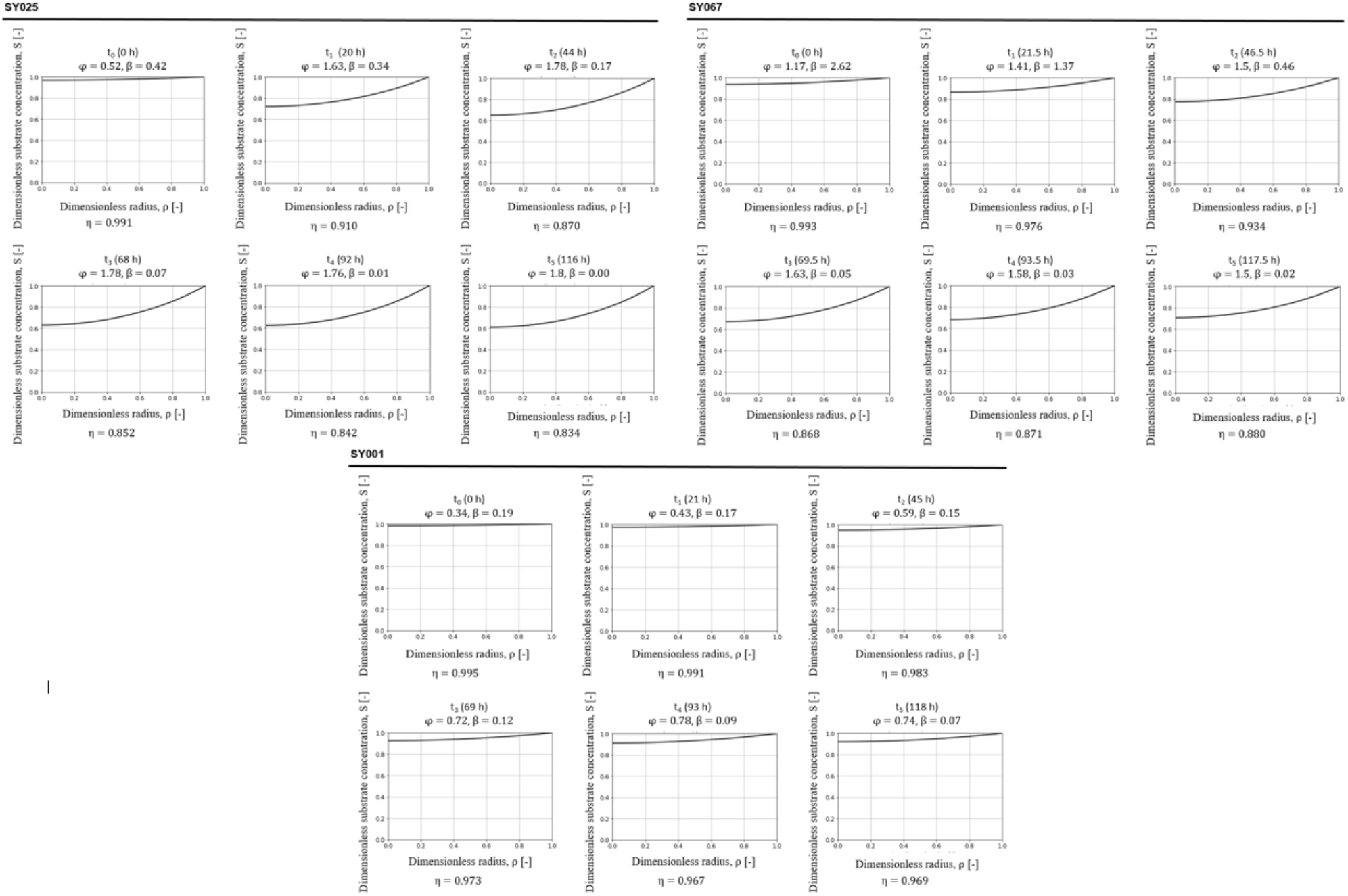
Estimated substrate concentration profiles inside the ImoYeast particle during beer fermentations for strains SY025, SY067 and SY001, in which: φ = Thiele modulus; η = effectiveness factor; and β = saturation parameter.

Based on the following criteria defined by Doran (2013): for φ < 0.3 then η ∼ 1 – not significant mass transfer limitations; and for φ > 3 then η << 1 – significant mass transfer limitations, mass transfer limitations are considered not to be the most significant phenomenon in the ImoYeast particle for all strains tested.

Results for SY025 strain point to the formation of a substrate concentration gradient inside the support during the process. At the end of the fermentation (116 h), 60% of bulk substrate concentration is available in the center of the particle. For strains SY067 and SY001, internal substrate concentrations were, respectively, 70% and 95% of those found in bulk liquid.

Scanning Electron Microscopy (SEM) was used to validate data from the reaction-diffusion model. Analysis revealed yeast accumulation inside the ImoYeast particles during fermentations carried out by all strains tested. It was not conclusive whether yeast cell concentration was higher at the center or the edge of the particles. It was also not possible to observe any distribution, orientation or positioning pattern for yeast accumulation inside the support. Fig. 6 presents micrographs obtained from ImoYeast containing strains SY025 and SY067.

**Fig. 6.**
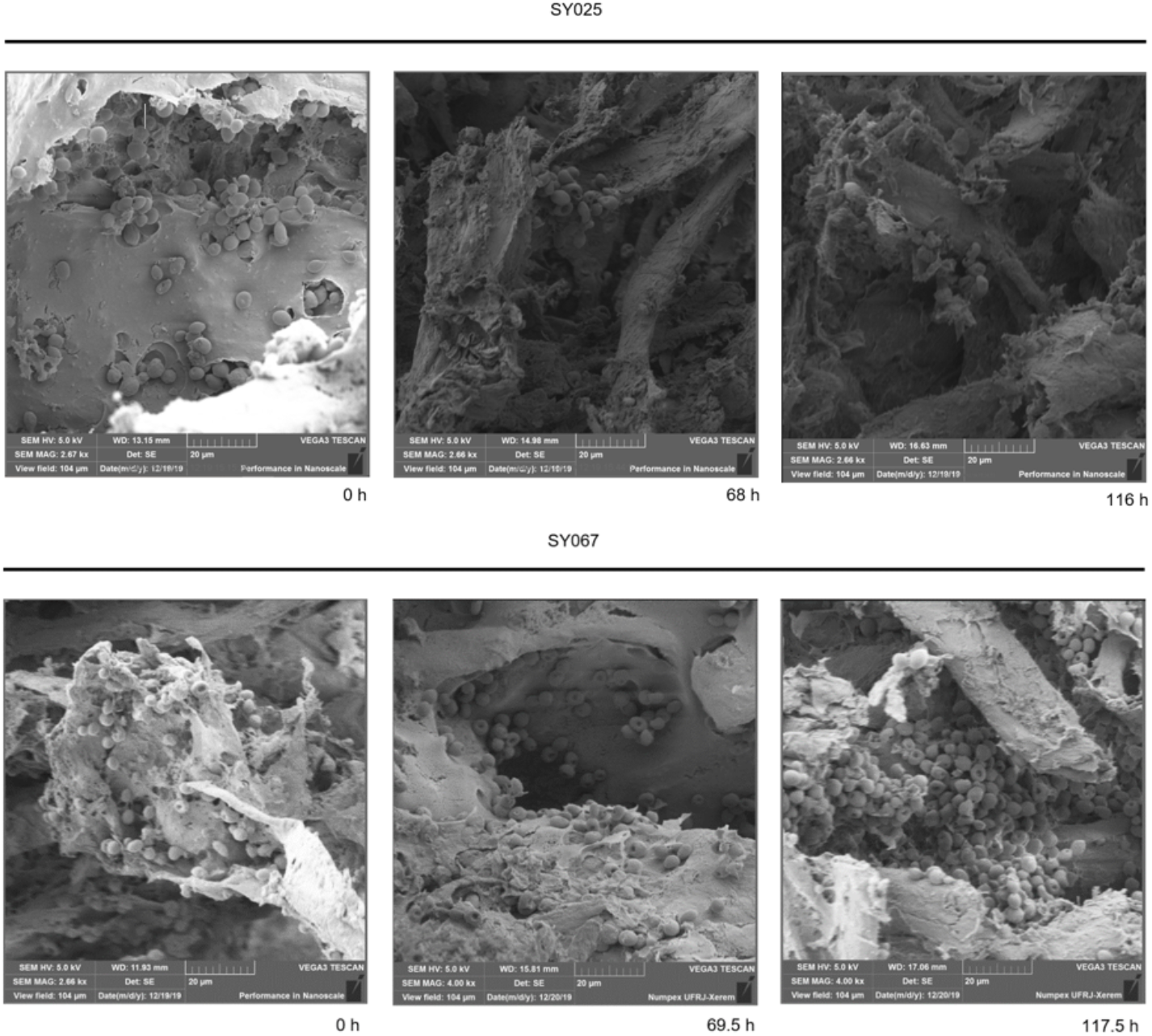
Scanning Electron Microscopy (SEM) of ImoYeast support during beer fermentations for strains SY025 (magnitudes of ca. 2670x) and SY067 (magnitudes of 2680x for 0 h and 4000x for 69.5 h and 117.5 h). Micrographs provide evidence of immobilized cells accumulation inside the support during the fermentation time.

### 3.3 Variation of initial biomass and substrate concentrations results in changes in flavor profile during fermentation

A central composite design experiment was performed using SY025 strain, to evaluate the response of immobilized cells to variations of initial biomass (X_0_) and substrate concentrations (S_0_), in respect to the formation of flavor-active compounds (Fig. 7).

**Fig. 7.**
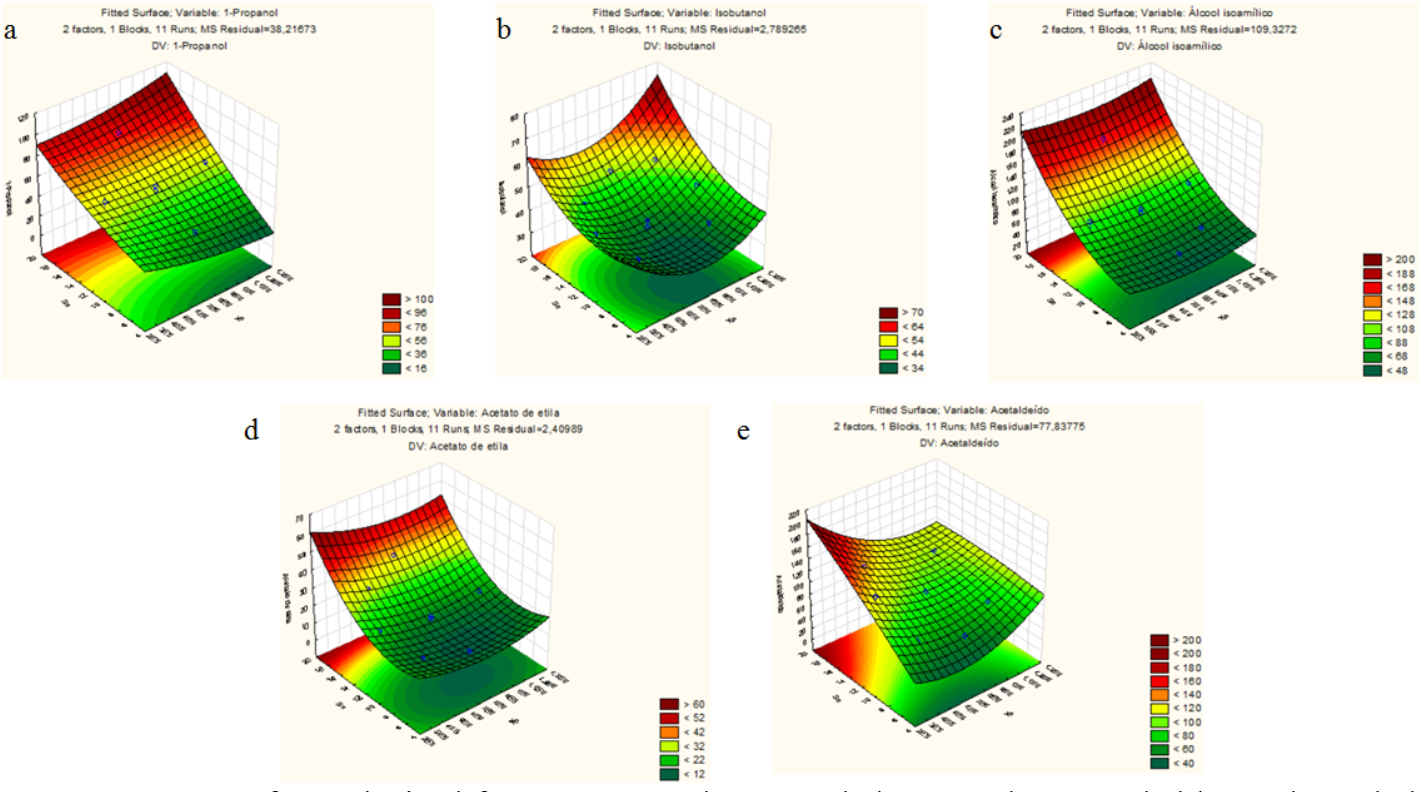
Response surfaces obtained for: **a** 1-propanol; **b** 2-methyl propanol; **c** 3-methyl butanol; **d** ethyl acetate; and **e** acetaldehyde, in beer fermentations with distinct biomass (X_0_) and substrate (S_0_) initial concentrations.

## 4. DISCUSSION

Immobilization efficiency was expected to decrease during fermentation, due to biomass growth on external regions of the particle, which favors cell release from the support (Tanaka et al. 1989). Yet, the immobilization method used was able to retain biomass according to the predefined criteria. This behavior is typical from methods of simultaneous infiltration and coating with gelation agents (Nedović et al. 2015; Verbelen et al. 2006), increasing cell density inside the support.

The ImoYeast support stimulates yeast cell growth during beer fermentation. For all investigated strains, Y_XS_ demonstrated a shift towards biomass formation by the immobilized cells in comparison to free cells. Considering the total biomass in the system, immobilized SY025 cells showed a 67% higher Y_XS_, compared to free cells. Similar results were obtained with immobilized SY067 and SY001 cells: 78% and 56% higher Y_XS_, respectively. The results match those described by Bezbradica et al. (2007), for the LentiKat^®^ support, which was described to stimulate cell growth.

Nevertheless, distinct growth patterns have been described for immobilized yeast, including higher, lower and equivalent biomass concentrations compared to free cell. The most accepted hypothesis to describe such variation is the formation of unique immobilization microenvironments depending on the support used. Usually, the higher the mass transfer limitation, the lower the cell concentration (Norton and D’Amore 1994).

Doran and Bailey (1986) showed that the substrate consumption rate by immobilized yeast cells during alcoholic fermentation was higher than that by free cells. Yet, the biomass growth rate was lower. Immobilization was then considered to affect biomass development due to cell-support and cell-cell physical contact and this results in morphological changes and alterations in the expression of cell wall proteins, like flocculins (Honigberg 2011), responsible for the formation of tridimensional yeast community structures. In the case of porous supports, high surface area is suitable for microbial development because of the available space cells have to interact with the matrix (Kregiel et al. 2012). This is a hypothesis to cell growth increasing in the ImoYeast support, a porous solid immobilization matrix.

An important aspect highlighted by Brányik et al. (2004) is the formation of an active-cell layer on the surface of the immobilization support, which favors cell release to bulk liquid. This is probably the case in the ImoYeast matrix. It is hypothesized that, once immobilized, yeast cells are stimulated to grow and those closest to the surface end up released to the liquid. Then, released cells remain active and keep growing now as free cells, which causes immobilized systems to show higher total biomass concentrations.

Immobilization efficiency (IE) values achieved during fermentation assays are directly related to the selected immobilization method. The results are strain dependent, based on biomass growth rate and concentration. Analyzing IE data and cell yield from substrate (Y_XS_), it is noticeable that biomass detachment from the support is proportional to biomass yield, corroborating the hypothesis from Brányik et al. (2004). An example is SY067 strain – immobilized cells showed higher total biomass growth, 1.78x the value from that of free cells. This strain exhibited the highest increase in biomass concentration, and immobilized SY067 cells presented also the lower IE value among the tested strains.

Johansen and Flink (1986) observed that, during alcoholic fermentations performed with immobilized *S. cerevisiae* in alginate particles, for alginate concentrations of 2-4%, the concentration of released cells to the medium was 2×10^5^ - 4×10^5^ cells/ mL. Using 1% alginate, cell detachment increased 100%, while for 5% alginate it was reduced in 80%. Therefore, since the alginate concentration used in the present study was 2%, cell detachment from ImoYeast support was expected.

Regarding main product formation, all three strains in suspension presented higher ethanol and lower glycerol yields in comparison to the immobilized cell configuration (Table 2). Free SY025 cells achieved higher ethanol yield in 9% than immobilized ones. Free SY067 and SY001 strains were superior to the immobilized configuration in 9% and 13%, respectively. About glycerol yields, immobilized SY025 cells achieved a 40% higher value than that of free cells, while immobilized SY067 and SY001 showed 53% and 19% higher values, respectively. Since the immobilized systems presented higher total biomass concentration, a higher glycerol formation was already expected, because of redox balance inside the cells, as described by van Dijken and Scheffers (1986).

*S. cerevisiae* immobilized in glass beads coated by gelatin showed higher specific substrate consumption rate (2x) and ethanol (45%) and glycerol production rates when compared to free cells (Doran and Bailey 1986). However, there were lower ethanol (free = 1.8±5% and immobilized = 1.4±17%) and glycerol (free = 0.11±3% and immobilized = 0.06±2%) yields from substrate. It was suggested that this occurred due to a metabolic shift from fermentation to storage of reserve polysaccharides (e.g., glycogen and trehalose), besides increased DNA and RNA intracellular concentrations. In accordance to concentration data, ethanol productivity was lower for immobilized cells, and there was a higher glycerol productivity. This is believed to be related to increased biomass concentration in the immobilized system (Brányik et al. 2004).

Regarding the increased acetaldehyde formation exhibited by immobilized SY025 cells, this is hypothesized to be due an increase on the substrate consumption rate (Doran and Bailey 1986). Once glucose consumption is stimulated, intracellular pyruvate concentration is increased and pyruvate formation becomes not only a metabolic control step, but also a bottleneck. Consequently, acetaldehyde concentration also increases. One part is excreted by the cells while the other is channeled into alcoholic fermentation. Finally, ethanol is used as a precursor of ethyl acetate by means of acetyl-CoA and acyl-CoA coenzymes (Galazzo and Bailey 1990).

The decreased formation of 2-methyl propanol and 3-methyl butanol by immobilized SY025 cells is considered to be related mainly to valine and leucine mass transfer limitations, important precursors on the production of these flavor compounds (Kobayashi et al. 2008). Another possibility is the excessive α-acetolactate excretion by immobilized yeast cells, another important precursor to both higher alcohols.

The increase in ethyl acetate formation by SY067 strain might be due to oxygen mass transfer limitations throughout the ImoYeast particle, resulting in the formation of this ester (Djordjević et al. 2016; Willaert and Nedovic 2006). However, this was the system that presented the lower immobilization efficiency. Then, even though oxygen mass transfer might have been limited, many cells were detached to the liquid medium. Thus, as the substrate consumption rate by immobilized cells was lower for this strain it is proposed that threonine, precursor to 1-propanol, is consumed before valine and leucine (precursors to 2-methyl propanol and 3-methyl butanol, respectively). This way, it is suggested that at the start of fermentation (in which IE values are higher), threonine mass transfer limitations are derisory, since its consumption is mainly carried out by the yeast active-layer on the matrix surface, resulting in 1-propanol formation. Finally, detached yeast cells that already consumed threonine would metabolize valine and leucine as the fermentation follows and IE decreases.

At last, for SY001 it is hypothesized that mass transfer limitations were at minimum and did not disturb sequential amino acid consumption, as was the case to the other strains investigated due to high cell detachment. It is also suggested that immobilization resulted in an increase of the substrate consumption rate, which made pyruvate formation an important metabolic bottleneck that channeled the cell metabolism into the production of 1-propanol, 2-methyl propanol and 3-methyl butanol. The higher concentration of ethyl acetate in the immobilized system can be explained by the activation of immobilized cell metabolism, besides oxygen exhaustion inside the ImoYeast matrix. Then, acetyl-CoA and acyl-CoA would be available for its synthesis.

Higher alcohol formation in immobilized yeast systems have been reported as being lower, equivalent or higher than those in free cell configurations (Djordjević et al. 2016). A direct comparison of the results obtained in this work to other is complex (Virkajärvi and Pohjala 2000; Willaert and Nedovic 2006), because of metabolic alterations that are hard to predict and understand, depending on the process configuration (e.g., batch or continuous), the immobilization method (e.g., adsorption or entrapment), the yeast strain (e.g., Ale or Lager), the support type (e.g., calcium alginate or porous glass), amongst other factors.

Willaert and Nedovic (2006) showed that yeast cells, immobilized to a support that stimulates biomass growth, produce higher alcohols in a greater extent, particularly 1-propanol, due to α-ketobutyrate pathway activation. Results from immobilized SY025 and SY001, but not from SY067, agree to this. On the other hand, a continuous brewing process with immobilized yeast exhibited increased acetaldehyde concentration (4x higher) and decreased concentrations of ethyl acetate, propanol and 3-methyl butanol (4x, 3x and 2x lower, respectively) in comparison to a traditional batch process with free cells (Brányik et al. 2008).

Few information is found in the literature concerning to the kinetics of cell release from immobilization supports. This is due to the variety of supports and immobilization methods available, besides all possible process conditions. The most common approach is to neglect this phenomenon in the studied system (Kostov et al. 2012). In this work, however, the immobilization efficiency (IE) parameter evidenced that cell release is significant and should not be neglected.

Therefore, a Michaelis-Menten/Monod type kinetics was proposed for cell release because of the tendency of IE to decrease along fermentation time and as cells grow. Thus, the specific cell release rate, k (h^-1^), was considered to be dependent on the immobilized cell biomass (m_Xim_).

The model residues from SY025 fermentation data were in accordance to similar studies with immobilized yeast found in literature: 0.419-14.8 (Kostov et al. 2012) and 0.07-0.81 (Parcunev et al. 2012).

In the sense of indicating that immobilization into the ImoYeast support stimulates biomass growth, model estimated parameters (especially μ_max_) for SY025 strain are in agreement with experimental data. Strong evidence of the influence the immobilization microenvironment exerts over microbial metabolism is the reduction of μ_max_ of released yeast cells in comparison to that of cells still attached to the matrix. The main hypothesis is that mass transfer limitations and cell-support interaction result in physiological responses towards metabolism activation (Brányik et al. 2005), and subsequently into higher yields of biomass and glycerol instead of ethanol. Considering Y_PXet_ and Y_PXgly_, the estimated values (Table 3) corroborate the experimental data. Free and immobilized cells present similar results for estimated Y_PXet_ (17.85 g _ethanol_/ g _free biomass_ and 16.47 g _ethanol_/ g _immobilized biomass_). Nevertheless, it is noticeable the likelihood of cell immobilization to result in lower ethanol formation. However, estimated Y_PXet_ for released cells is drastically higher (181.47 g _ethanol_/ g _released biomass_). For estimated Y_PXgly_, the results are similar: free cells (0.57 g _glycerol_/ g _free biomass_) produce less glycerol than immobilized ones (0.85 g _glycerol_/ g _immobilized biomass_). Yet, when released to the liquid medium, their metabolism is displaced towards ethanol formation, since Y_PXgly_ becomes 0.08 g _glycerol_/ g _released biomass_. Therefore, the modeling of SY025 cell fermentation indicate that once in direct contact to the ImoYeast support, their metabolism is channeled into biomass, glycerol and flavor-active compounds formation (Brányik et al. 2004; Kourkoutas et al. 2004; Norton and D’Amore 1994).

The modeling of SY067 indicates a reduction in the specific maximum growth rate (μ_max_) when cells are immobilized (0.14 h^-1^ free cells to 0.093 h^-1^ immobilized), which is even more significant when cells are released from the immobilization support (0.016 h^-1^). Using a 13% malt wort, Parcunev et al. (2012) also demonstrated that top-fermenting immobilized yeasts showed lower μ_max_ than those in suspension (0.016 h^-1^ free cells to 0.009 h^-1^ immobilized). At first, it could be hypothesized the arise of a substrate gradient with significant impact over substrate availability inside the support, hindering biomass growth (Brányik et al. 2005; Doran and Bailey 1986). But the reaction-diffusion modeling estimated the substrate gradient in the ImoYeast particle not to be limiting. Therefore, the alternative hypothesis is that SY067 is sensible to the effects from high cell density and interaction between cell wall and solid support. Quorum sensing phenomena are possibly involved in cell growth rate reduction. In other words, due to cell-cell and cell-solid interactions, SY067 metabolism is believed to channel into the formation of structural and energetic reserve polysaccharides, instead of accelerating cell growth rate inside the ImoYeast support.

Yet, as previously stated, cells located at the surface of the support ended up being released from it (Brányik et al. 2004) and continued to grow This phenomenon is explained by Karel et al. (1985), showing that in some cases cells adsorbed to a solid can either grow or release their daughter-cells to the liquid medium..

Estimated ethanol (Y_PXet_) and glycerol (Y_PXgly_) yields from biomass corroborates to experimental data (see Tables 2 and 3). Similar results were obtained for SY001 strain, to which estimated μ_max_ values suggest its immobilization to ImoYeast support increases cell growth rates (0.08 h^-1^ free cells to 0.47 h^-1^ immobilized). It is also in agreement with the results from a bottom-fermenting yeast strain in a 13% malt wort (Parcunev et al. 2012) that showed higher μ_max_ for immobilized (0.0125 h^-1^) than free (0.009 h^-1^) cells. The explanation relies on the high cell concentration within the control volume, which accelerates reaction rates. At last, estimated ethanol and glycerol yields presented the same trend of experimental data: immobilized cells (Y_PXet_ = 17.04 g _ethanol_/ g _immobilized biomass_) produced less ethanol than free cells (Y_PXet_ = 17.53 g _ethanol_/ g _free biomass_) and more glycerol (immobilized: Y_PXgly_ = 0.88 g _glycerol_/ g _immobilized biomass_; and free: Y_PXgly_ = 0.82 g _glycerol_/ g _free biomass_).

With respect to the reaction-diffusion modeling, Thiele modulus (φ) and effectiveness factor (η) were chosen for quantitative evaluation of the ImoYeast support performance, since they are engineering parameters commonly used for evaluating the efficiency of heterogeneous reaction systems (Weisz 1973). Yet, literature rarely reports such analyzes applied to immobilized cell systems and the majority of models neglect substrate diffusion effects. Brányik et al. (2005), however, discuss that substrate mass transfer limitations are the most obvious hypothesis to explain physiological alterations in fermentation processes. Therefore, as ImoYeast is a new immobilization support, it was important to investigate whether mass transfer limitations would be significant or not.

In an overview, it is highly difficult to define whether the observed physiological alterations are only due to cell-support interaction or also to the formation of an immobilization microenvironment by substrate diffusion limitations, or even some interaction between both these factors.

In the SY025 case, physiological modifications might be in part related to the formation of a substrate gradient inside the particle. During the fermentation, values of η decreased from 0.991 (0 h) to 0.834 (116 h), indicating a decrease of the substrate diffusion efficiency through the matrix. Values of φ also indicate that, ranging from 0.52 at 0 h to 1.8 at 116 h. The same tendency was observed for SY067 and SY001 strains. In other words, as the fermentation evolves, the support efficiency decreases, exacerbating the impacts of substrate gradient (Pilkington et al. 1998).

For SY067 cells, η values changed from 0.993 to 0.880 and φ values from 1.17 to 1.5, from the start to the end of the fermentation, respectively. SY001 strain showed η values from 0.995 to 0.969 and φ values from 0.34 to 0.74.

Since estimated mass transfer limitations were minimal, we considered that the observed physiological alterations were mainly caused by cell-cell and cell-support interactions.

Scanning Electron Microscopy (SEM) images obtained along the fermentation assays showed that yeast cells were able to colonize the interior of the ImoYeast particle, not only at its surface (Fig. 6). Even with the formation of a substrate gradient inside the support, cells thrived and were able to multiply. Micrographs show interactions between cells and of cells with the support, indicating growth as colonies adsorbed to the internal walls of the matrix. This agrees with the formation of tridimensional structures of microbial communities due to flocculins expression (Honigberg 2011). Yet, there was no easily recognizable pattern of biomass growth inside the solid because of heterogeneous development of cell colonies (Walsh and Malone 1995).

Regarding the variation of biomass (X_0_) and substrate (S_0_) initial concentrations, it was observed for strain SY025 that, none of the conditions tested resulted in the overproduction of 1-propanol and 2-methyl propanol, in concentrations above their threshold limits (800 mg/L and 200 mg/L, respectively). Thus, considering such compounds, SY025 immobilization does not negatively interfere in final product quality and aroma profile. On the other hand, all experimental conditions resulted in acetaldehyde concentrations above the threshold limit (25 mg/L), which is still a challenge to be addressed for ImoYeast-based immobilized systems since acetaldehyde excess might confer beer with a not balanced green leaves aroma (Olaniran et al. 2017).

## STATEMENTS AND DECLARATIONS

### Authors’ contributions

All authors conceived and designed research. TA conducted experiments. MC performed scanning electron microscopy analysis. MC, MB, BB and TB contributed new reagents or analytical tools. All authors analyzed data. TA wrote the manuscript. MB, BB and TB revised the manuscript. All authors read and approved the manuscript.

## Acknowledgments

We thank the technical support provided by Rayane Gonçalves Pereira da Silva in scanning electron microscopy analysis.

## LIST OF SYMBOLS

μ = specific cell growth rate (h^-1^)

m_X_ = cell mass (g _biomass_)

q_S_ = specific substrate consumption rate (g _substrate_/ (g _biomass_. h))

q_Pet_ = specific ethanol formation rate (g _ethanol_/ (g _biomass_. h))

q_Pgly_ = specific glycerol formation rate (g _glycerol_/ (g _biomass_. h)

μ_im_ = specific immobilized cell growth rate (h^-1^)

m_Xim_ = immobilized cell mass (g _immobilized biomass_)

k = cell release specific rate (h^-1^)

μ_rel_ = specific released cell growth rate (h^-1^)

m_Xrel_ = released cell mass (g _released biomass_);

q_S, im_ = specific substrate consumption rate by immobilized cells (g _substrate_/ g _immobilized biomass_. h)

q_S, rel_ = specific substrate consumption rate by released cells (g _substrate_/ g _released biomass_. h)

q_Pet, im_ = specific ethanol formation rate by immobilized cells (g _ethanol_/ g _immobilized biomass_. h)

q_Pet, rel_ = specific ethanol formation rate by released cells (g _ethanol_/ g _released biomass_. h)

q_Pgly, im_ = specific glycerol formation rate by immobilized cells (g _glycerol_/ g _immobilized biomass_. h)

q_Pgly, rel_ = specific glycerol formation rate by released cells (g _glycerol_/ g _released biomass_. h)

μ_max_ = maximum specific growth rate (h^-1^) m_S_ = substrate mass (g _substrate_)

K_SX_ = saturation Monod constant (g _substrate_)

Y_XS_ = biomass yield from substrate (g _biomass_/ g _substrate_)

Y_PXet_ = ethanol yield from biomass (g _ethanol_/ g _biomass_)

Y_PXgly_ = glycerol yield from biomass (g _glycerol_/ g _biomass_)

μ_max, im_ = maximum specific immobilized cells growth rate (h^-1^) m_S_ = substrate mass (g _substrate_)

K_SX, im_ = immobilized cells saturation Monod constant (g _substrate_)

Y_XS, im_ = immobilized biomass yield from substrate (g _biomass_/ g _substrate_)

Y_PXet, im_ = ethanol yield from immobilized biomass (g _ethanol_/ g _biomass_)

Y_PXgly, im_ = glycerol yield from immobilized biomass (g _glycerol_/ g _biomass_)

μ_max, rel_ = maximum specific released cells growth rate (h^-1^)

K_SX, rel_ = released cells saturation Monod constant (g _substrate_)

Y_XS, rel_ = released biomass yield from substrate (g _biomass_/ g _substrate_)

Y_PXet, rel_ = ethanol yield from released biomass (g _ethanol_/ g _biomass_)

Y_PXgly, rel_ = glycerol yield from released biomass (g _glycerol_/ g _biomass_)

k_max_ = maximum specific cell release rate (h^-1^)

m_Xim_ = immobilized cell mass (g)

k_m_ = immobilized cell mass in which release rate is half the maximum (k = k_max_/ 2)

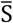 = dimensionless substrate concentration (-)

S_SURF_ = substrate concentration at the surface of the particle (g _substrate_/ L)

*ρ* = dimensionless radius (-) φ = Thiele modulus (-)

D_eff_ = substrate effective diffusivity through immobilization support (m^2^/ h)

β = saturation parameter (-)

